# Evaluating nongenetic maternal transmission and post-weaning persistence of gut microbiota in cross-fostering rabbits

**DOI:** 10.1101/2025.07.30.667589

**Authors:** Shi-Yi Chen, Hancheng Gao, Junkun Zhou, Mingyan Cui, Xinyang Tian, Wenqiang Sun, Song-Jia Lai, Yinghe Qin, Xianbo Jia

## Abstract

Gut microbiota is widely recognized as an important source contributing to phenotypic variation of diverse host traits in livestock. In this regarding, it is promising to improve offspring production traits through direct selection on parental individuals with the desirable gut microbiota, and this is termed microbiome breeding. However, the extent of nongenetic maternal transmission of gut microbiota and its post-weaning persistence remains poorly understood. To fill this gap, we cross-fostered 120 newborn rabbits that are offspring of 20 does in this study, and compared gut microbiota of all kits at the weaning (35 days of age) and two weeks post-weaning, respectively. Based on variance component analysis, we found that the large proportions of phenotypic variation of four alpha diversity metrics for kits at weaning can be explained by the nursing does, ranging from 0.218 ± 0.106 for Evenness index to 0.464 ± 0.125 for the number of observed features. The significant effect of nursing does was supported further by comparing three beta diversity metrics among different relationships, using Kruskal-Wallis test and Dunn’s post-hoc comparisons. In contrast, the biological does showed no significant effect on gut microbiota composition of pre-weaning kits. In post-weaning kits, the influence of nursing does on gut microbiota was not obviously decreased, despite there was increasing contributions of additive genetic effects. In conclusion, this study provides direct evidence in rabbits that offspring gut microbiota is predominantly shaped by the nongenetic maternal transmission, with these maternal influences persisting post-weaning. These results indicate the possibility that the direct selection on parental gut microbiota will alter offspring gut microbiota.

## Introduction

In livestock, gut microbiota is extremely complex and has been recognized to strongly interact with diverse physiological functions of host, such as diet digestion and nutrient absorption, mucosal immune development, and homeostasis maintenance (Hanning and Diaz-Sanchez, 2015; Holman and Gzyl, 2019). The dramatically reduced costs in high-throughput DNA sequencing have enabled large-scale profiling of gut microbiota, hence facilitating its growing use in exploring associations with host phenotypes related to feed efficiency, growth, health, and other important traits in various livestock species (e.g., Pérez-Enciso et al., 2021; He et al., 2022; Tian et al., 2024). Therefore, it appears promising to influence individual phenotypes through deliberately altering the gut microbiota (Clemmons et al., 2019; Upadhaya and Kim, 2022; Larzul et al., 2024).

There are different approaches that can be employed for manipulating the gut microbiota composition in animals, including antibiotic administration, nutritional interference, cross-fostering, cohousing, and microbial transfer (Ericsson and Franklin, 2015). In the context of livestock breeding, however, it is desirable to select the parental individuals with or without gut microbiota information, and consequently influence gut microbiota composition of the reproduced offspring. Many studies have revealed that there are host genetic effects on gut microbiota composition in livestock (e.g., Wallace et al., 2019; Ishida et al., 2020; Qin et al., 2022; Wang et al., 2023; Lecoeur et al., 2024), and hence means it is feasible to indirectly alter offspring microbiota composition via genetically selecting the parents (Casto-Rebollo et al., 2023; Larzul et al., 2024). However, the potential benefits of such indirect selection may be limited, because few heritable microbial taxa have been identified and most of them exhibit the relatively low heritability. Therefore, it would be valuable to know whether direct selection on the gut microbiota of parental individuals could lead to predictable alterations among their offspring; this selection process is also termed microbiome breeding (Mueller and Linksvayer, 2022). If the selected superiority on microbiota could be, at least partially, transmitted persistently to offspring, gut microbiota composition may be considered a pseudo-heritable trait and subject to selection-based changes across generations.

A critical consideration when directly selecting gut microbiota composition among parental individuals is its transgenerational fidelity, as this fundamentally determines offspring responses to selection (Mueller and Linksvayer, 2022). In mammals, maternal gut microbiota can be vertically transmitted to offspring through multiple pathways, including the prenatal exposure via the placenta, perinatal inoculation during birth, and postnatal colonization via the breastfeeding and physical contacts (Bogaert et al., 2023; Zhuang et al., 2024). Furthermore, in the polytocous mammals, gut microbiota can be horizontally transmitted among littermates through the shared environmental exposure and social interactions (Wanelik et al., 2023). Due to these transmission pathways, individual and population-level gut microbiota compositions could be maintained for ten generations found in inbreeding mouse lines (Moeller et al., 2018). However, the observed maternal transmission of gut microbiota may be partially confounded by the maternal additive genetic effects, which is hard to be distinguished. To our knowledge, maternal nongenetic transmission and persistence of gut microbiota have not been specifically studied yet for the polytocous livestock. To fill this gap, we established a cross-fostering population of domestic rabbits (*Oryctolagus cuniculus*) in this study, aiming to evaluate maternal transmission of gut microbiota to suckling kits and post-weaning persistence. Our findings will provide crucial insights into the application of microbiome breeding in polytocous livestock.

## Materials and methods

### Study design and sampling

The cross-fostering experiment was illustrated in Figure 1. A total of 30 multiparous does of New Zealand White rabbits were initially selected, and all of them did not have the known genetic relationships with each other during the previous generations. After subject to the light-assisted synchronization procedure of ovulation, these does were naturally mated with different bucks at the same day. Between two does that kindling at the same day, about half of newborn kits were randomly selected and cross-fostered with each other. All kits were weaned at 35 d (days of age), after when each kit was individually housed with ad libitum consumption of a commercial pellet diet until 49 d. Other feeding and management practices followed our standard procedures.

**Figure 1.**
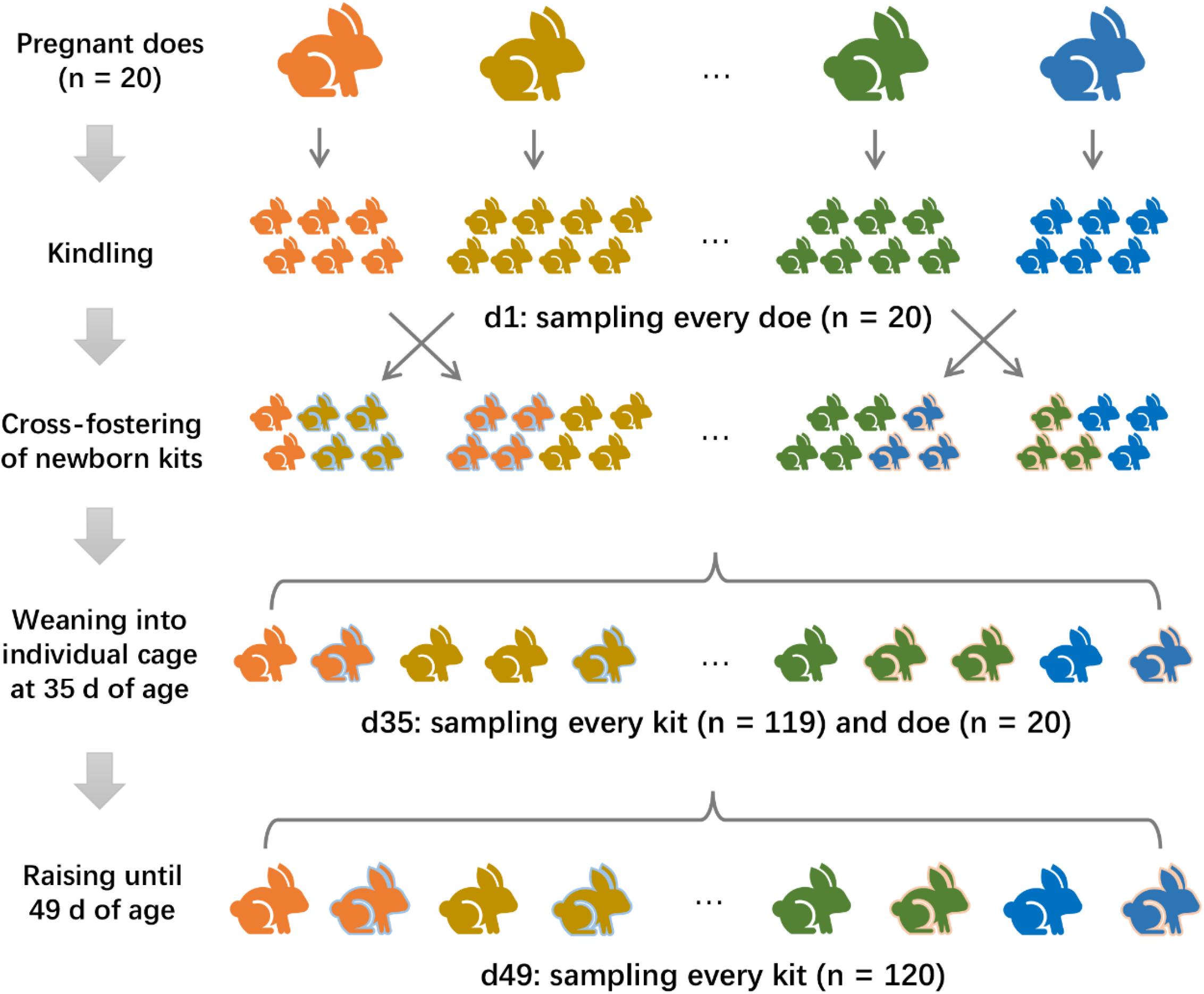
Design of cross-fostering experiment and sample collection.

A total of 120 kits from 20 does that had at least five healthy offspring at the end of the experiment were finally included for analyses. The rectal feces samples were collected individually for the 20 does at the kindling day (1 d) and weaning day (35 d), and rectal feces samples were also collected individually for the 120 kits at 35 d and 45 d. As the feces sample of one kit collected at 35 d was destroyed during the storage in liquid nitrogen, there were 279 feces samples in total used for sequencing microbial 16S rRNA gene. Other individual information, including the gender and live body weight, was not recorded.

### 16S rRNA gene sequencing and bioinformatic analyses

Bacterial DNA was extracted using QIAamp BiOstic Bacteremia DNA Kit (Qiagen, Shanghai, China), and the concentration and quality were determined by NanoVue Plus (GE Healthcare, Piscataway, USA). Using the universal primers (338F: 5’-ACT CCT ACG GGA GGC AGC AG-3’ and 806R: 5’-GTG GAC TAC HVG GGT WTC TAA-3’), we amplified V3-V4 region of 16S rRNA gene with HOTSTAR Taq Plus Master Mix Kit (Qiagen, Shanghai, China), using the procedure of an initial denaturation step at 95 °C for 4 min, 20 cycles of 95 °C for 1 min, 56 °C for 45 sec and 72 °C for 1 min, and an extension step at 72 °C for 7 min using a Bio-Rad CFX96 thermal cycler (Bio-Rad, Hercules, USA). After purification using QIAquick PCR Purification Kit (Qiagen, Shanghai, China), amplicons with a total amount of ≥ 3 μg and OD260/280 ratio ≥ 1.8 were used to prepare sequencing libraries using Illumina DNA Sample Preparation Kit (Illumina, San Diego, USA) according to official instruction. Finally, the libraries were sequenced on Illumina HiSeq™ 2000 platform for generating 300 bp paired-end reads. We performed the bioinformatic analyses of 16S rRNA sequencing data on the QIIME2 platform v2024.10 (Bolyen et al., 2019). In brief, raw paired-end reads were first merged using VSEARCH software v2.22.1 (Rognes et al., 2016), to which at most three mismatches and minimum length of 20 bp were allowed for the overlapped region. Low-quality sequences were discarded if they contained any 5-bp window with an average Q-score < 30 (Bokulich et al., 2013). The denoising and clustering algorithm in Deblur software v1.1.1 (Amir et al., 2017) was used for removing both artificial and chimera sequences, and then inferring the amplicon sequence variants (ASVs). Based on the rarefaction curves generated by “diversity alpha-rarefaction” function (Supplementary Figure S1), every sample was randomly rarefied to the minimum number of 28,023 sequences for avoiding the bias of different sequencing depths across samples (i.e., no sample was excluded in this dataset). The sequences of ASVs were taxonomically assigned using the pre-fitted sklearn-based taxonomy classifier and Greengenes database v202409 (DeSantis et al., 2006). Using the function of “diversity core-metrics-phylogenetic”, we calculated four alpha diversity metrics of Shannon index, the number of observed features, Faith’s phylogenetic diversity (Faith’s PD), and Evenness index for every sample, as well as three beta diversity metrics of Jaccard distance, Bray curtis distance, and weighted unifrac distance.

### Statistical analyses

After discarding the low abundance ASVs that are present in less than 28 individuals (i.e., 10% of total samples), square root-normalized relative abundances were obtained to visualize overall relationships among all samples using multiscale PHATE software v0.3.0 and default parameters (Kuchroo et al., 2022). For alpha diversity metrics, we first statistically compared the differences among the four groups of both does and kits at the three ages (i.e., does at 1 d and 35 d, and kits at 35 d and 49 d), using the non-parametric and rank-based Kruskal-Wallis test, together with Dunn’s post-hoc comparisons, provided in rstatix R package v0.7.2 (Kassambara, 2023). We additionally compared the changed alpha diversity metrics of kits between 35 d and 49 d within every nursing doe.

Due to the full-sib structure of kits in this study, we further estimated individual additive genetic effects and common litter environmental effects on individual alpha diversity metrics sampled at 35 d or 49 d, using the linear mixed effect model as follow:

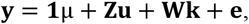

where **y** is the phenotype vector of alpha diversity metric of kits. **1** is a vector of ones and μ is the overall mean. **u** and **k** are the random effects of individual additive genetic and common litter environmental effects, with the assumption of **u**∼N(0,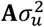) and **k**∼N(0,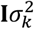), respectively. Here, **A** is additive genetic similarity matrix for this full-sib population, together with its variance of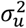; **I** and 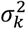 are the identity matrix and common litter environmental variance, respectively. **e** is a vector of residual effects with **e**∼N(0,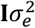). All variance components were estimated using the average-information REML method implemented in the blupf90 software (Masuda, 2018; Misztal et al., 2018). The proportions of phenotypic variance explained by individual additive genetic effects (*h*^2^) and common litter environmental effects (*k*^2^) were obtained as 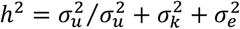 and 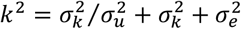, respectively.

For beta diversity metrics of kits sampled at 35 d or 49 d, we statistically compared their means among four relationship groups, including: (1) two kits with same biological and nursing does (bDoe/nDoe, 338 pairs), (2) two kits with different biological but same nursing does (−/nDoe, 340 pairs), (3) two kits with same biological but different nursing does (bDoe/-, 335 pairs), and (4) two kits with different biological and nursing does (−/-, 13,208 pairs). To avoid the extremely unequal sample sizes, 338 pairs (i.e., the mean of other three groups) were randomly sampled from the fourth group. As above, Kruskal-Wallis test and Dunn’s post-hoc comparisons were used.

## Results

### Gut microbiota composition

A total of 29.3 million raw paired-end reads were obtained, with an average of 104,954 pairs per sample. After quality filtering, we retained an average of 50,203 sequences per sample, ranging from 28,023 to 82,131. There were 8,185 ASVs in the rarefied abundance table, and 55%, 90%, and 93% of them could be taxonomically annotated at the genus, family, and order levels, respectively. The top three dominant phylum were Bacteroidota, Bacillota, and Verrucomicrobiota, respectively; and the compositions are comparable among the four groups of does and kits at the three ages (Supplementary Figure S2). Based on 1,718 ASVs that were present in more than 10% of total samples, clustering patterns of samples are shown in Figure 2 according to three top dimensions, which revealed that all samples of both does and kits at the three ages were evenly distributed on the whole.

**Figure 2.**
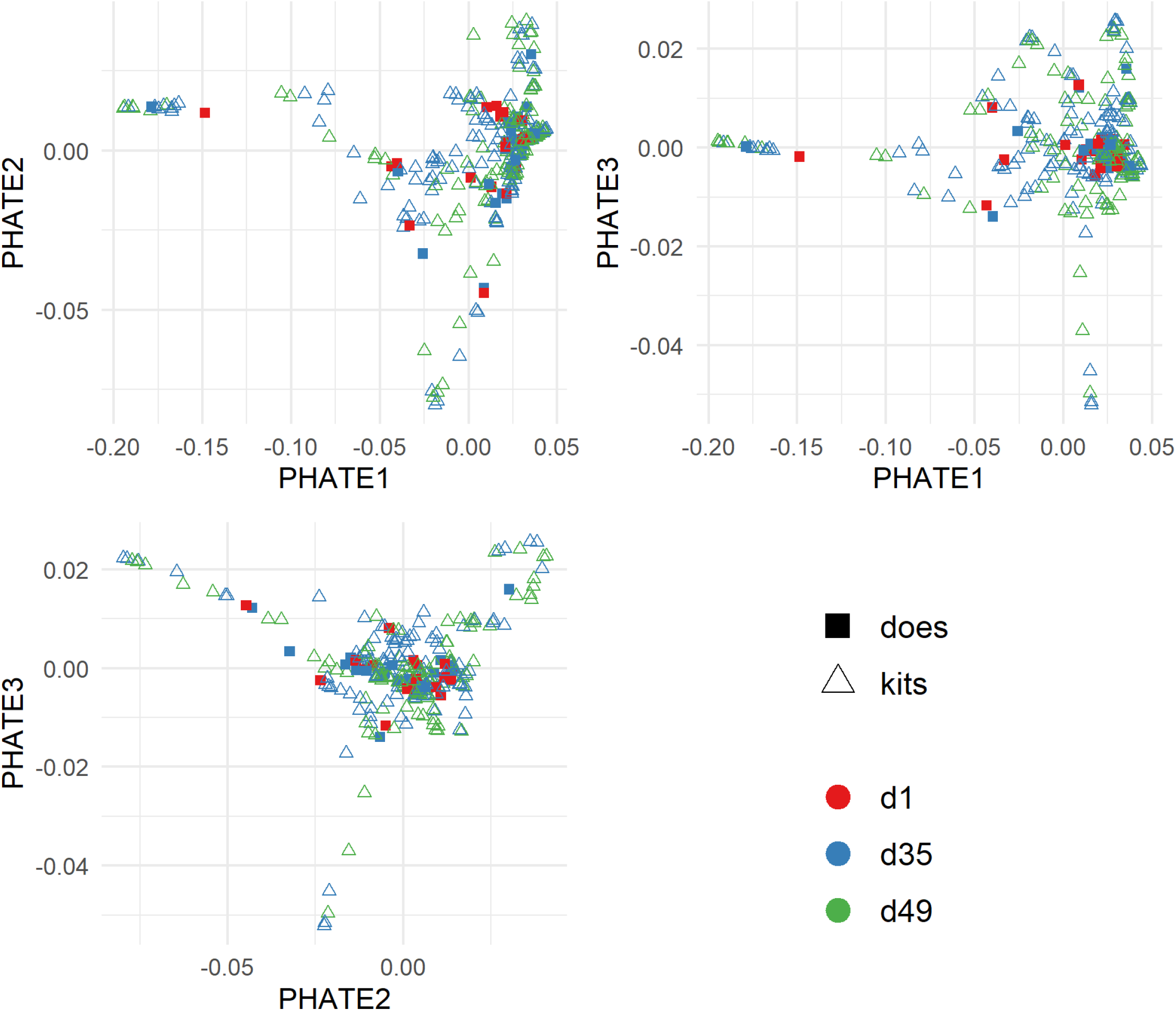
Multiscale PHATE visualizations of microbiota composition among all samples.

### Maternal transmission

With the exception of Evenness index, three metrics of alpha diversity were differed significantly among the four groups of both does and kits at the three ages (Figure 3; Kruskal-Wallis test *p* < 0.01). Pairwise comparisons revealed that the highest alpha diversity was observed for the does and there was no significant difference between the kindling day and weaning day (Dunn test *p* > 0.05). The weaning kits had the lowest alpha diversity, with significant differences with that of does at 1 d or 35 d (Dunn test *p* < 0.05). Post-weaning, alpha diversity of kits significantly bounced back at the 49 d (Dunn test *p* < 0.05), but still slightly lower than that of does. The same trends were observed for the Evenness index, although their *p* values did not reach statistically significant level. Within every nursing doe, the overall increased alpha diversity was observed post-weaning (Supplementary Figure S3).

**Figure 3.**
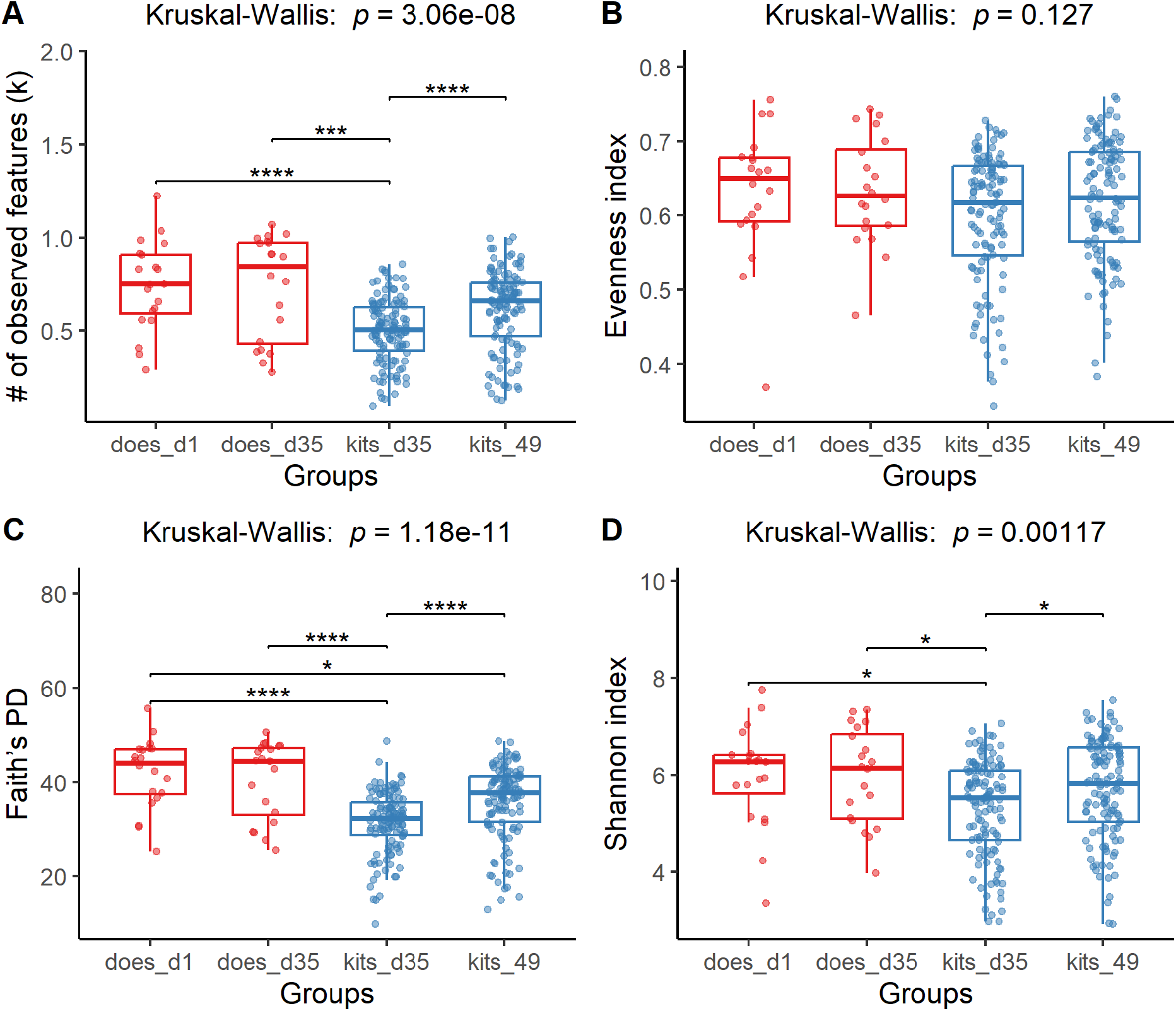
Statistical comparisons of four alpha diversity metrics among the four groups of both does and kits at three age stages. Faith’s PD: Faith’s phylogenetic diversity. The *p* value of Kruskal-Wallis test is shown to the top of boxplots, followed by Dunn’s post-hoc comparisons (* *p* < 0.05, ** *p* < 0.01, *** *p* < 0.001, and **** *p* < 0.0001).

Based on variance component analysis of alpha diversity metrics (Table 1; mean ± standard error, SE), we found that the large proportions of phenotypic variation of kits at weaning can be explained by the nursing does (i.e., the litter environmental effect) for the number of observed features (0.469 ± 0.126), Evenness index (0.218 ± 0.106), Faith’s PD (0.464 ± 0.125), and Shannon index (0.345 ± 0.120), respectively. However, the estimated individual additive genetic effect did not deviate from zero, while the models converged successfully only for the number of observed features (0.017 ± 0.095). Based on three beta diversity metrics, we statistically compared the pairwise similarities of kits at weaning among the four groups of different relationships (Figure 4A). These results revealed that their pairwise similarities were mainly determined by whether the two kits had same nursing does or not; i.e., there were significant influences of nursing does on the pairwise similarities (Dunn test *p* < 0.01), whereas the biological does did not have the significant effects on them (Dunn test *p* > 0.05).

**Table 1.** Estimates (SE) of variance components and proportions for four alpha diversity metrics

| Metrics | Age | Variance components<sup>1</sup> |  |  | Proportions<sup>2</sup> |  |
| --- | --- | --- | --- | --- | --- | --- |
| | | $\sigma_u^2$ | $\sigma_k^2$ | $\sigma_e^2$ | $h^2$ | $k^2$ |
| # of observed features | d35 | 487 (2714) | 15086 (5862) | 15236 (3002) | 0.017 (0.095) | 0.469 (0.126) |
|  | d49 | 8950 (7445) | 22839 (9346) | 15916 (5174) | 0.193 (0.167) | 0.458 (0.138) |
| Evenness index | d35 | \ | 17.46 (9.17) | 59.54 (8.47) | \ | 0.218 (0.106) |
|  | d49 | 24.30 (16.61) | 8.40 (7.10) | 31.93 (11.08) | 0.367 (0.243) | 0.125 (0.110) |
| Faith's PD | d35 | \ | 21.91 (8.53) | 23.36 (3.33) | \ | 0.464 (0.125) |
|  | d49 | 11.66 (10.13) | 28.69 (11.99) | 23.75 (7.26) | 0.186 (0.167) | 0.428 (0.137) |
| Shannon index | d35 | \ | 0.36 (0.16) | 0.65 (0.09) | \ | 0.345 (0.120) |
|  | d49 | 0.24 (0.22) | 0.30 (0.16) | 0.52 (0.16) | 0.230 (0.210) | 0.270 (0.133) |

<sup>1</sup> $\sigma_u^2$ : additive genetic variance; $\sigma_k^2$ : common litter environmental variance; $\sigma_e^2$ : residual variance. The slash (\) indicates additive genetic effects were not included because the model did not converge successfully.
<sup>2</sup>The explained proportions of phenotypic variance by additive genetic effects ( $h^2$ ) and common litter environmental effects ( $k^2$ ).

**Figure 4.**
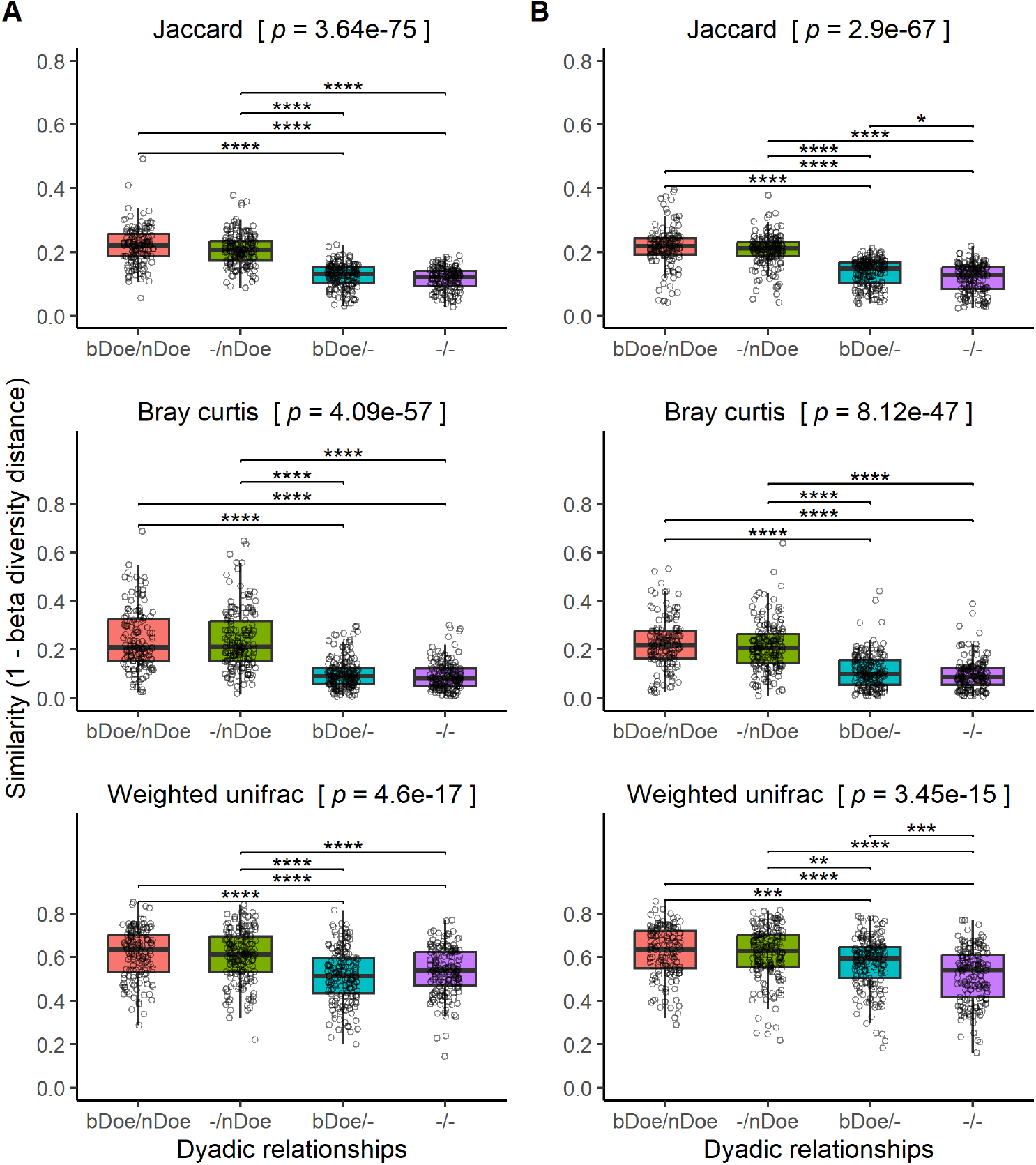
Statistical comparisons of three beta diversity metrics among four different relationships of kits at 35 d (A) or 49 d (B) of age. bDoe/nDoe: with same biological and nursing does, -/nDoe: with different biological but same nursing does, bDoe/-: with same biological but different nursing does, -/-: with different biological and nursing does. The *p* value of Kruskal-Wallis test is shown in the square bracket, followed by Dunn’s post-hoc comparisons (* *p* < 0.05, ** *p* < 0.01, *** *p* < 0.001, and **** *p* < 0.0001).

### Post-weaning persistence

There were still significant effects of nursing does on phenotypic variation of alpha diversity for kits at 49 d, only with the slightly reduced proportions that were explained by the litter environmental effect (Table 1). However, we detected individual additive genetic effects on the four alpha diversity metrics at 49 d, having the estimates of heritability ranging from 0.186 ± 0.167 to 0.367 ± 0.243, although with the large SE. Similarly, the significant effects of nursing does on pairwise similarities of kits were remained at 49 d (Dunn test *p* < 0.01), according to the statistical comparisons of beta diversity metrics (Figure 4B). We further calculated the paired similarities between 35 d and 49 d for all kits (n = 119), and their similarities were comparable with that of nursing siblings based on the three beta diversity metrics (Table 2). When regressing individual alpha diversity of kits at 49 d on that at 35 d (Figure 5), the statistically significant positive relationships (*p* < 0.05) were revealed for the four metrics although the regression coefficients varied from 0.055 to 0.535.

**Table 2.** Paired similarities of kits between 35 d and 49 d of age based on beta diversity

| **Metrics** | **Mean** | **Median** | **SD** | **Min** | **Max** |
| --- | --- | --- | --- | --- | --- |
| Jaccard | 0.215 | 0.209 | 0.048 | 0.101 | 0.408 |
| Bray curtis | 0.217 | 0.197 | 0.111 | 0.042 | 0.579 |
| Weighted unifrac | 0.600 | 0.621 | 0.119 | 0.266 | 0.833 |

**Figure 5.**
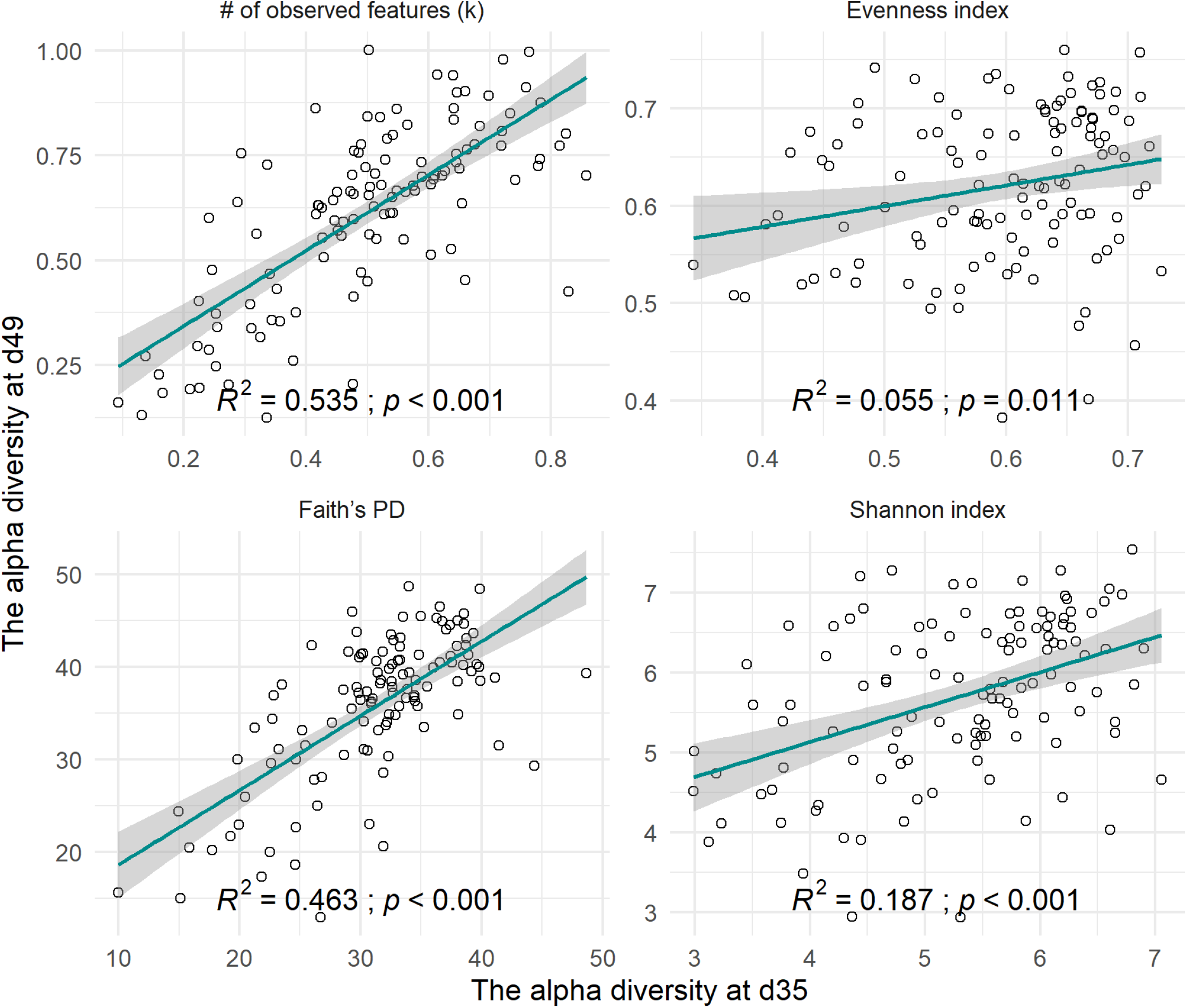
Line regression of individual alpha diversity of kits at 49 d on that at 35 d of age. Faith’s PD: Faith’s phylogenetic diversity. The regression line and 95% confidence intervals are shown by green line and grey area.

## Discussion

Rabbits are the most recently domesticated livestock in southern France ∼1400 years ago (Irving-Pease et al., 2018). Owe to the high reproductive efficiency with short gestation period and large litter sizes, each doe can reproduce over 50 rabbits marketed per year under the optimized conditions (Huneau-Salaün et al., 2015), yielding more than 60 kg of high-quality meat (Kumar et al., 2025). Another merit of rabbits is the ability to effectively utilize plant fiber fractions that are indigestible to human, and this process is largely facilitated by gut microbiota (Velasco-Galilea et al., 2018). Therefore, rabbits have been popularly raised in Asian and European countries for producing high-quality meat. As many studies have identified the associations between gut microbiota composition and host health and production phenotypes in rabbits (e.g., Chen et al., 2019; Fang et al., 2020; Velasco-Galilea et al., 2021; Liu et al., 2022; Tian et al., 2024), it may be possible to improve offspring phenotypes of interest through selecting the gut microbiota among parental individuals. To address this concern, we quantified the nongenetic maternal transmission of gut microbiota to suckling rabbits and assessed its persistence post-weaning in this study.

For pre-weaning traits in livestock, individual direct genetic effects and maternal or litter common environmental effects are often confounded together (Bijma, 2006), and the specific experimental designs and sophisticated statistical models are required to distinguish them. In this study, we cross-fostered newborn kits between the paired does to separate the confounded individual direct additive genetic effects on gut microbiota composition; such cross-fostering approach was recently used for dissecting the effect of litter breed composition on the growth, survival, and health traits of rabbits (Bigot et al., 2024). Based on the partition of phenotypic variance of alpha diversity and statistical comparisons of beta diversity, our results clearly revealed that there was no individual additive genetic effect on gut microbiota composition of kits at weaning, whereas the individual genetic effect was present at two weeks post-weaning. The most obvious post-weaning change in individual gut microbiota observed in this study was the significantly increased diversity, which was previously reported in growth rabbits (Fang et al., 2020). Accordingly, we propose that host genetic influences on the gut microbiota in rabbits are varied pre-and post-weaning, likely related to the increasing microbiota diversity. There may be another cause that the relaxed impacts of common litter environment on gut microbiota among post-weaning kits would also result into the increased individual direct genetic effects.

In contrast to individual direct genetic effects, our results demonstrated that gut microbiota composition of kits at weaning is significantly influenced by the nursing does, hence indicating the effective nongenetic maternal transmission of gut microbiota to offspring. Using the cow-to-calf model, Zhuang et al. (2024) similarly found the significant vertical transmission of gut microbiota from mother to offspring in dairy cattle. Through feeding probiotics to pregnant mothers in mice, rats, and pigs, maternal transmission of gut microbiota to offspring was directly observed, contributing to the compositional assemblages among offspring (Buddington et al., 2010). Because rabbits are polytocous, however, mutual horizontal transmission of gut microbiota would be present among littermates through the shared environment and social interactions during the suckling period, which will further promote their resembles with nursing does. In pigs, about 30% and 13% of piglet gut microbiota could be source-tracked to their mothers at 7 d after birth and post-weaning, respectively; and the cage mates contributed 54% to the gut microbiota at 10 d post-weaning (Tancredi et al., 2025). Collectively, these studies provide direct evidence that maternal gut microbiota could be effectively transmitted to offspring via both vertical and horizontal transmission during the sucking period, especially for the polytocous livestock. Due to the significant transmission to offspring, therefore, maternal gut microbiota may be considered as a pseudo-heritable trait.

In rabbits, as well as other livestock, gut microbiota composition considerably changes among different ages, especially immediately after the weaning (Fang et al., 2020; Fu et al., 2024; Biada et al., 2025). During the first one to two weeks post-weaning, rabbit gut microbiota composition undergoes rapid reorganization in response to dietary transition and environmental changes, potentially causing the increased mortality rates during this critical period (Rashwan and Marai, 2000). In this study, therefore, we further investigated gut microbiota composition of kits at two weeks post-weaning to quantify the post-weaning persistence of maternal influences. Our results confidently revealed that the significant influences of nursing does on offspring’s gut microbiota composition at two weeks post-weaning remained almost comparable to the effects observed at weaning. The similar results were observed in pigs (Tancredi et al., 2025), which revealed the majority of maternally transmitted microbes pre-weaning could persist post-weaning. While only the limited studies found in literature have addressed this topic to date, available evidence could demonstrate robust post-weaning persistence of maternally transmitted gut microbiota in both rabbits and pigs.

Two limitations of this study are acknowledged. First, individual growth traits, such as average daily weight gain post-weaning, were not measured among offspring, hence precluding the identification of maternally transmitted microbes associated with individual phenotypes. If it is intended to improve offspring production performance through direct selection on parental gut microbiota, it is critical to verify whether the altered gut microbiota among offspring, as observed in this study, could result into desirable phenotypic outcomes. Second, we don’t know whether sex differences exist in maternally transmitted gut microbiota among offspring because individual sex was not recorded. As only weak effect of sex on gut microbiota composition was observed in rabbits (Ye et al., 2021), this issue may be of limited importance, especially regarding the nongenetic maternal transmission of gut microbiota.

## Conclusion

In this study, we experimentally established a cross-fostering population in rabbits and quantified the nongenetic maternal transmission of gut microbiota and post-weaning persistence. Our results clearly demonstrated that the nursing does significantly shape offspring gut microbiota composition, and the nursing-based transmission dominates over individual direct genetic effects. Furthermore, there are robust post-weaning persistence of the maternally transmitted gut microbiota. These results provide direct evidence supporting the possibility to implement microbiome breeding in rabbits.

## Acknowledgments

This study was financially supported by Earmarked Fund for China Agriculture Research System (CARS-43-A-2), and Science & Technology Department of Sichuan Province (2021YFYZ0033).

## Ethics statement

The animal study was approved by Institutional Animal Care and Use Committee of Sichuan Agricultural University. The study was conducted in accordance with the local legislation and institutional requirements.

## Software and data repository resources

All the data supporting the results of this study are included in the article and in the Additional file.

