## Supplementary Figures for "Evaluating nongenetic maternal transmission and post-weaning persistence of gut microbiota in cross-fostering rabbits"

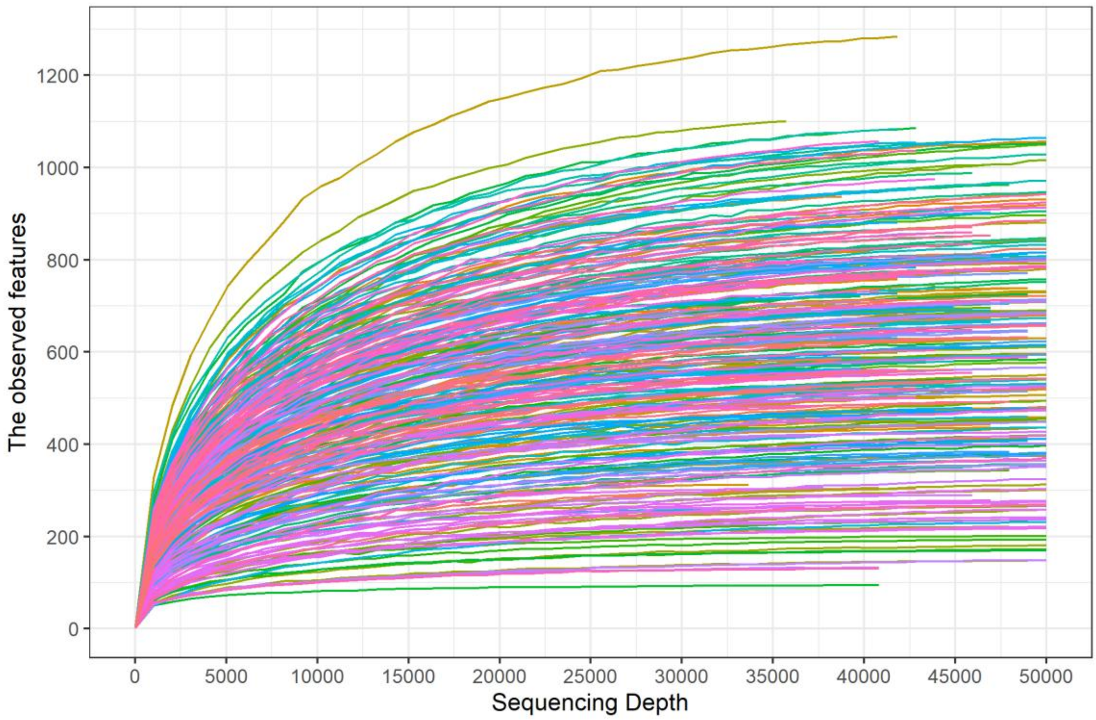


**Supplementary Figure S1. Rarefaction curves of the number of observed features among different sequencing depth**.

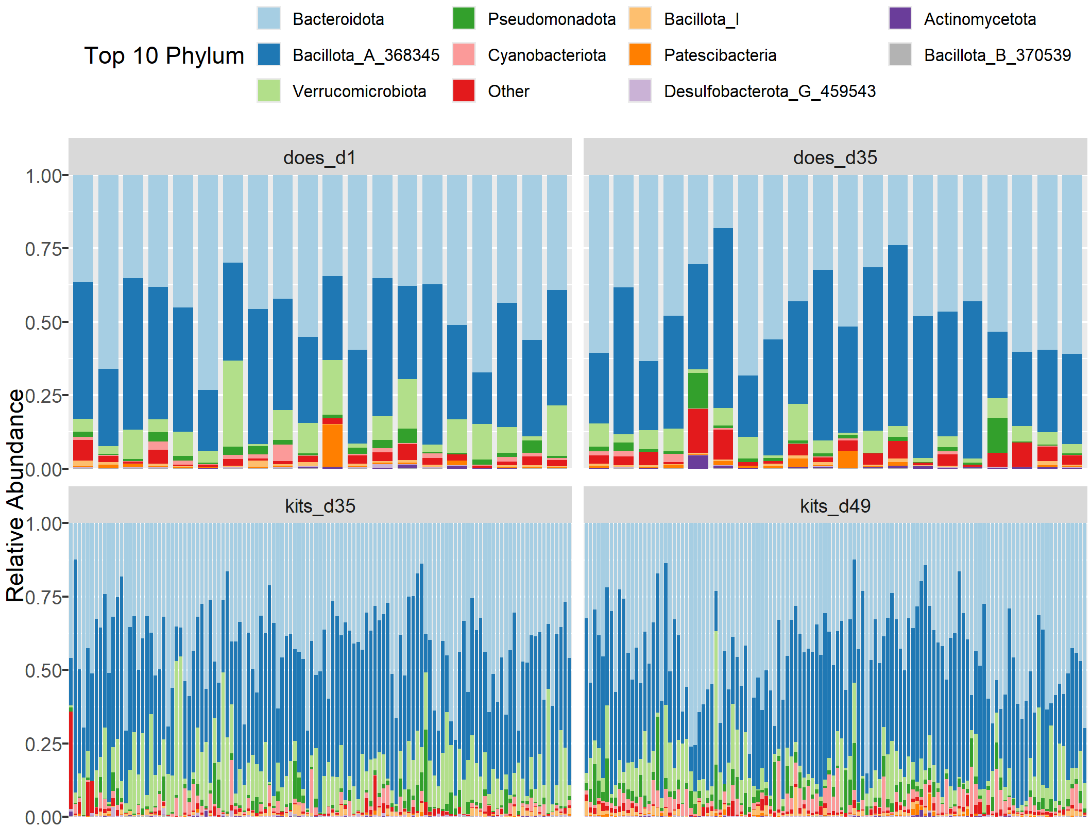


**Supplementary Figure S2. Taxonomical compositions at the phylum level for both does and kits at the three ages.**

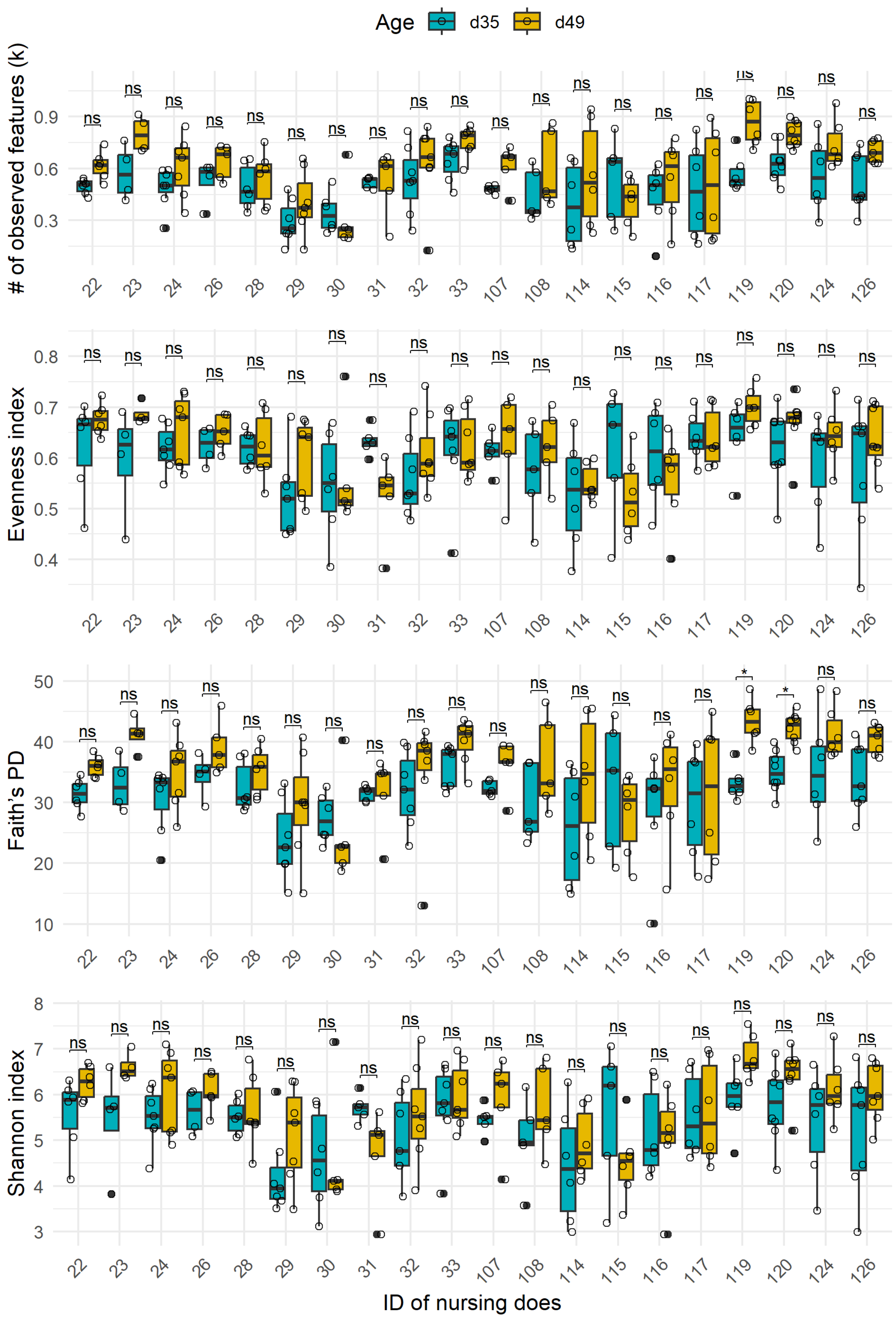


**Supplementary Figure S3. Comparisons of individual alpha diversity of kits between 35 d and 49 d of age within every nursing doe**. Faith's PD: Faith's phylogenetic diversity. The Kruskal-Wallis test is applied (ns: $p > 0.05$, *: $p < 0.05$).
